# In-silico identification of archaeal DNA-binding proteins

**DOI:** 10.1101/2024.08.09.607351

**Authors:** Linus Donvil, Joëlle A.J. Housmans, Eveline Peeters, Wim Vranken, Gabriele Orlando

## Abstract

The rapid advancement of next-generation sequencing technologies has generated an immense volume of genetic data. However, this data is unevenly distributed, with well-studied organisms being disproportionately represented, while other organisms, such as from archaea, remain significantly underexplored. The study of archaea is particularly challenging due to the extreme environments they inhabit and the difficulties associated with culturing them in the laboratory. Despite these challenges, archaea likely represent a crucial evolutionary link between eukaryotic and prokaryotic organisms, and their investigation could shed light on the early stages of life on Earth. Yet, a significant portion of archaeal proteins are annotated with limited or inaccurate information.

Among the various classes of archaeal proteins, DNA-binding proteins are of particular importance. While they represent a large portion of every known proteome, their identification in archaea is complicated by the substantial evolutionary divergence between archaeal and the other better studied organisms.

To address the challenges of identifying DNA-binding proteins in archaea, we developed Xenusia, a neural network-based tool capable of screening entire archaeal proteomes to identify DNA-binding proteins. Xenusia has proven effective across diverse datasets, including metagenomics data, successfully identifying novel DNA-binding proteins, with experimental validation of its predictions.

Xenusia is available as a PyPI package, with source code accessible at https://github.com/grogdrinker/xenusia, and as a Google Colab web server application at https://colab.research.google.com/drive/1c4eb4sEz8OsBqHL62XDFrqmwa7CxImww?usp=sharing.

## INTRODUCTION

The rise of next generation sequencing technologies has brought us in a time where a lot of genetic information is being generated. This information, together with annotations such as domains and functions, is continuously added to big databases like UniProt ^1^, which stores all the details about protein sequences. UniProt has grown significantly, now holding over 250 million sequences and still growing. However, upon closer examination, it becomes evident that there is an uneven distribution of information among various organisms. Widely studied organisms like *Homo sapiens, Escherichia coli* and *Danio rerio* (zebrafish) are well-represented, while other species suffer from neglectance, resulting in an underrepresentation. This is what experts call an “observational selection bias”, where most species end up with limited data and mainly inaccurate annotations, pushed to the sidelines without much detailed information ^2^.

Currently, there is a big scientific debate about our ability to predict the fold of completely new proteins, especially when dealing with proteins that have very limited or no evolutionary information available. Even AlphaFold ^3^, which in recent years revolutionized structural biology, struggles to deal with proteins that have limited evolutionary information ^4^. A large part of the archaeal proteins belong to this group. By August 2024, UniProt contained only 3932 archaeal entries with evidence of existence at the protein level. The main problem includes the complete lack of information for a large portion of archaeal organisms. Recent studies on metagenomics data ^5,6^ highlighted the inability of many archaeal species that live in extreme environments to grow *in vitro*. Fortunately, new sequencing technologies have been successful in extracting genomic and proteomic information from such extremophiles. While some of these proteins showed homology with both bacteria and eukaryotes, others were completely novel ^6^. The lack of homologues makes novel proteins extremely challenging to annotate, requiring ad hoc bioinformatics approaches to assign their putative function. Given the extreme environments that many of these organisms live in, they likely have folds that we have never seen before, and for which there are almost no similar proteins available for comparison. Investigating these unknown areas could greatly improve our knowledge about protein sequence and structural space.

This exploration is not only intellectually stimulating but also holds practical implications. The extreme environments where archaea thrive suggest that their proteins may possess unique molecular structures with potential applications. Their resilience to non-canonical conditions could offer valuable insights for industrial processes that demand protein stability under extreme circumstances.

A particularly important group of proteins are those that have the ability to interact with DNA. These proteins make up around 2-5% ^7^ of a species’ proteome and are essential for various biological processes, including gene regulation and DNA repair ^8,9^. While our current knowledge of archaeal protein structures is limited, the archaeal DNA-binding domains appear quite similar to those in bacteria, concerning structure and topology ^10^. However, the evolutionary gap is so vast that traditional tools, which rely on sequence similarity, often struggle to identify these regions. To address this challenge, we created a neural network-based tool called Xenusia. This tool operates without the need for multiple sequence alignment (MSA). The exclusion of MSA-based features has two significant effects: firstly, Xenusia provides extremely fast predictions, allowing proteome-wide analyses. Secondly, the lack of reliance on evolutionary features enhances Xenusia’s resistance to overfitting while simultaneously allowing it to identify convergent evolution. We demonstrate its effectiveness on diverse datasets, including metagenomics data. Xenusia successfully identifies novel DNA-binding proteins, a finding confirmed through experimental examination. Xenusia is available as a PyPi package and as source code at https://github.com/grogdrinker/xenusia, and as a Google Colab web server application at https://colab.research.google.com/drive/1c4eb4sEz8OsBqHL62XDFrqmwa7CxImww?usp=sharing.

## RESULTS

### COMPUTATIONAL APPROACH

The main challenge for annotating archaeal proteins is the prominent lack of data, hindering the efficacy of machine learning training directly on archaeal DNA-binding proteins. In the development of our tool, we consciously avoided possible prediction biases or overfitting, therefore exclusively using bacterial proteins for training (and parameter optimization) and reserving archaeal proteins for validation.

#### FEATURE ENCODING

As previously discussed, a primary challenge in this task is mitigating overfitting. A common approach to address this issue is by reducing the number of model parameters. To achieve this, we opted against utilizing parameter-intensive architectures based on embeddings. Instead, we employed a feature encoding strategy based on biophysical predictions. Specifically, we utilized single sequence-based predictors, using the services developed by Kagami and coauthors ^11^, to encode the input protein sequences into feature vectors. Detailed information on the feature encoding process can be found in the Methods section.

#### PREDICTION PIPELINE

Xenusia is a neural network-based tool that implements a two-step prediction pipeline, where each step is carried out by a distinct neural network, designated as NN1 and NN2, respectively. The first network (NN1) receives the initial feature vector as input and is trained to predict the residues directly interacting with DNA (refer to the contact dataset in the Methods section). The primary objective of NN1 is to identify the biophysical characteristics of amino acids that interact with DNA.

The second network (NN2) is trained on a dataset of DNA-binding domains (refer to the domain dataset in the Methods section). NN2 takes the output of NN1, comprising putative DNA-binding residues, as input and predicts which residues belong to DNA-binding domains. This architecture enables the model to learn how the patterns of potentially DNA-interacting residues (predicted by NN1) contribute to the formation of a DNA-binding domain (predicted by NN2).

The rationale behind this design is closely linked to overfitting concerns. Extracting domain-specific information from a limited set of proteins lacking detectable evolutionary relationships is challenging. By incorporating intermediate tasks, such as predicting DNA-interacting residues, the network is better constrained, thereby reducing the risk of overfitting.

NN1 utilizes a recurrent neural network (RNN) architecture, which encodes information from the entire protein sequence. In contrast, NN2 employs a sliding window-based feed-forward neural network, focusing on localized sequence features.

#### FINAL SCORE

The final score for a protein, which estimates the likelihood that an archaeal protein interacts with DNA, is determined by taking the maximum value of a sliding window average over the residue-based predictions generated by NN2. This approach is based on the observation that a protein’s DNA-binding capability is often attributed not to the entire protein, but rather to specific domains within it.

### COMPUTATIONAL VALIDATION

The performance of NN1, identifying DNA-binding residues, was validated by a 5-fold cross-validation on the contact dataset (see Methods section), resulting in an area under the ROC curve (AUC) of 0.75, with an average precision (AP) of 0.245. Given the low performance, particularly in terms of the AP, this model cannot provide reliable information to experimentalists that are interested in, for example, mutating potential DNA-binding residues. However, once NN1 is connected to NN2, the performances of Xenusia are significantly boosted. Figure 1 shows the increased performances in predicting archaeal DNA-binding domains (NN2) when including DNA-binding residue information (NN1). This highlights the importance of integrating information from various sources for reliable predictions.

**Figure 1:**
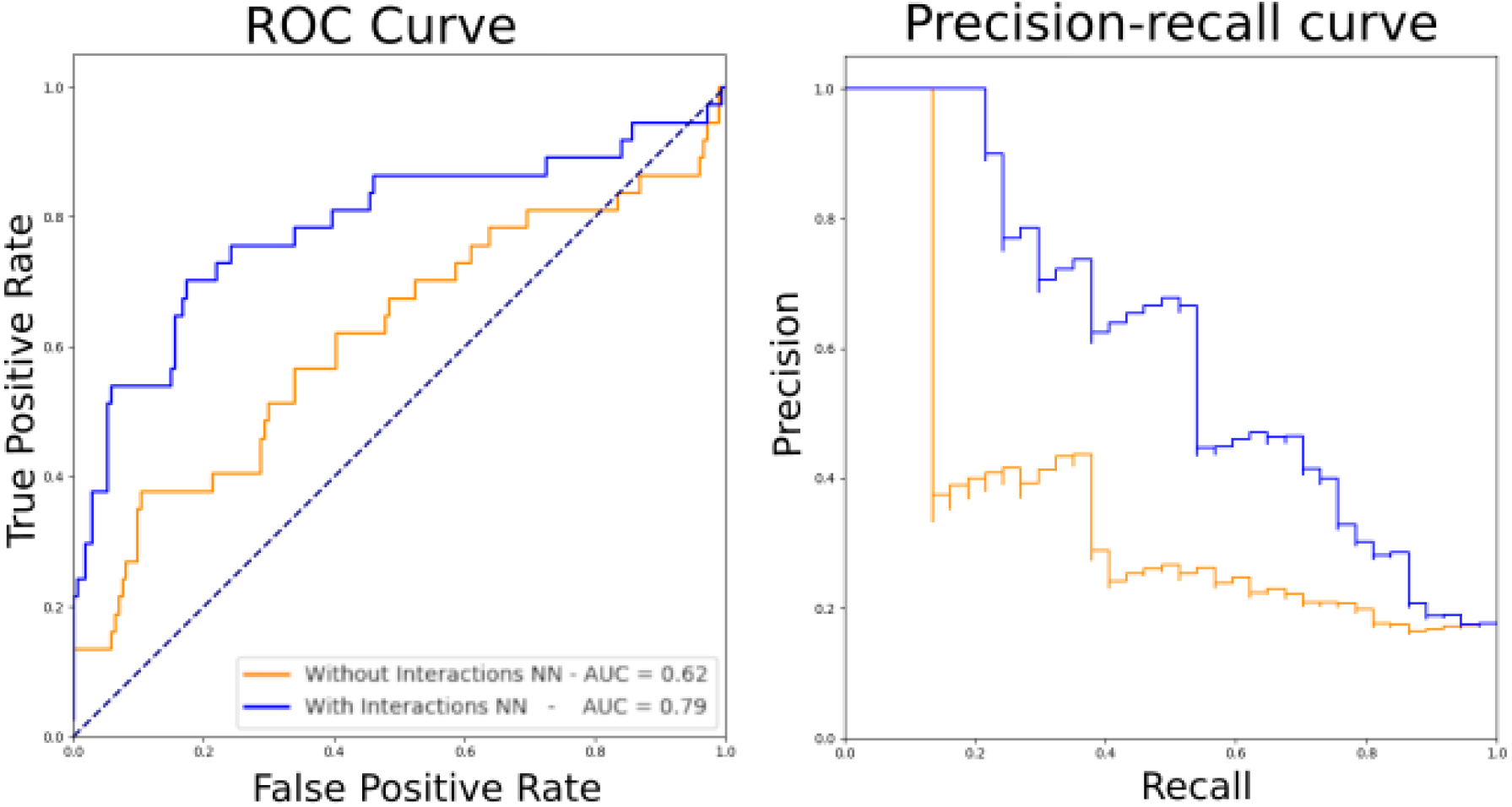
Performance of Xenusia. The tool’s performance on the validation dataset, composed of archaeal proteins only, is compared for a neural network without (orange) and with (blue) the inclusion of an intermediate prediction on interaction residues. Left, ROC curve; right, precision-recall curve.

The final performances of Xenusia result in an AUC of 0.79 and 0.60 for the ROC and Precision-Recall curve, respectively. In Table 1, the performance of Xenusia is compared to state-of-the-art methods. As the tool’s main purpose is to prioritize archaeal DNA-binding proteins, we had to select a threshold and compile a confusion matrix, enabling fair comparison with state-of-the-art binary classifiers. The threshold was set equal to the specificity value of the best performing state-of-the-art method and the same approach was performed for the predictor called local-DPP, as it also provides probabilities.

**Table 1:** Performance comparison of Xenusia with state-of-the-art methods for identifying DNA-binding proteins. Since Xenusia outputs a probability score, a threshold was chosen for binarization, allowing comparison with state-of-the-art binary predictors. The threshold was set to equal the specificity value of the best performing state-of-the-art method in our validation procedure (IDNA-Prot). AUC, area under the curve; MCC, Matthew’s correlation coefficient.

| Method | Accuracy | Sensitivity | Specificity | AUC | MCC |
| --- | --- | --- | --- | --- | --- |
| Xenusia | 77.73 | 70.27 | 79.31 | 0.79 | 0.414 |
| IDNA-Prot | 76.08 | 60.00 | 79.31 | - | 0.330 |
| DNAbinder | 63.51 | 56.76 | 64.94 | - | 0.169 |
| PseDNA-Pro | 64.93 | 72.97 | 63.22 | - | 0.278 |
| Local-DPP | 70.62 | 27.03 | 79.88 | 0.59 | 0.064 |

Table 1 shows that Xenusia provides the best predictions compared to all other available methods. The Matthew’s correlation coefficient (MCC), recognized as the best measure to summarize a confusion matrix, surpasses the second best predictor, IDNA-Prot with 0.330, by 25%. It is important to note that most available methods lack quality scoring information, making them unsuitable for proteome-wide functional studies based on experiments.

### LARGE SCALE ANALYSIS OF ASGARD METAGENOMICS

Asgard is a proposed superphylum of archaea that has only recently been discovered ^12^. These uncultivated archaea show intermediate characteristics between eukaryotes and prokaryotes. They have been found in samples coming from the Arctic sea ^13^ and other extreme environments, such as acid hot-springs ^12^. Homology analyses performed on a subclass of the Lokiarchaeota superphylum revealed that the protein phylogeny of these organisms is very uncommon: about one third of the proteins is completely new, while two thirds could be linked to other archaeal, bacterial or eukaryotic proteins ^14^. This portion of completely new proteins is enormous, making standard homology-based computational annotation tools unsuitable for the analysis of this taxonomic group. Since these organisms live in the Arctic sea or hot springs, their proteins have likely evolved to maximize functionality in extreme environments. This means that functional annotation of their proteins could offer new insights into the relationship between sequence and structure. Such information could be useful in specific industrial processes. We ran Xenusia on a dataset consisting of 16,946 Asgard proteins. The 10 best scoring proteins are reported in Supplementary Table 1. The first hit, OLS30781, is a homologue of a bacterial metalloprotease. This class includes enzymes with various functions, i.e. DNA repair ^15^. Interestingly, one of their distant homologues was crystallized in a complex with DNA (PDB ID: 1V14). Additionally, it is worth mentioning that ABC transporters are also commonly linked to DNA repair processes ^16^. The second and third hit, OLS30429 and OLS21705 respectively, are characterized proteins that are annotated as helix-turn-helix (HTH)-motif transcription regulators ^17,18^. These two proteins have relatively distant homologues, sharing only 46% and 37% sequence identities with their closest non-hypothetical relatives. The fifth and sixth hits, OLS17998 and OLS16705 respectively, are also homologues of archaeal HTH transcription factors. Regarding the protein homologue of the seventh hit, OLS19303 or also known as the response regulator SaeR, several studies have been performed on the equivalent protein of Staphylococcus aureus, defining the presence of a DNA-binding domain that acts as transcription regulator ^19^. Interestingly, the ninth highest-scoring protein, OLS30097, is an unannotated sequence, lacking any known annotated homologues. Despite this, the AlphaFold predicted structure reveals a distinct HTH structure. In Figure 2, a comparison between this model and the crystal structure of an archaeal HTH DNA-binding protein (5box, chain A:1:111) is shown. Notably, the unannotated Asgard protein displays a reversed topology compared to the crystal structure: the former begins with a globular domain and concludes with a long alpha helix, while the latter starts with a long alpha helix and finishes with the globular domain. This suggests independent, convergent evolution. It also clarifies why standard alignment algorithms and even structural alignment tools struggle to detect similarities between these two proteins. Supplementary Figure 1 shows the result of the structural alignment of the two proteins.

**Figure 2:**
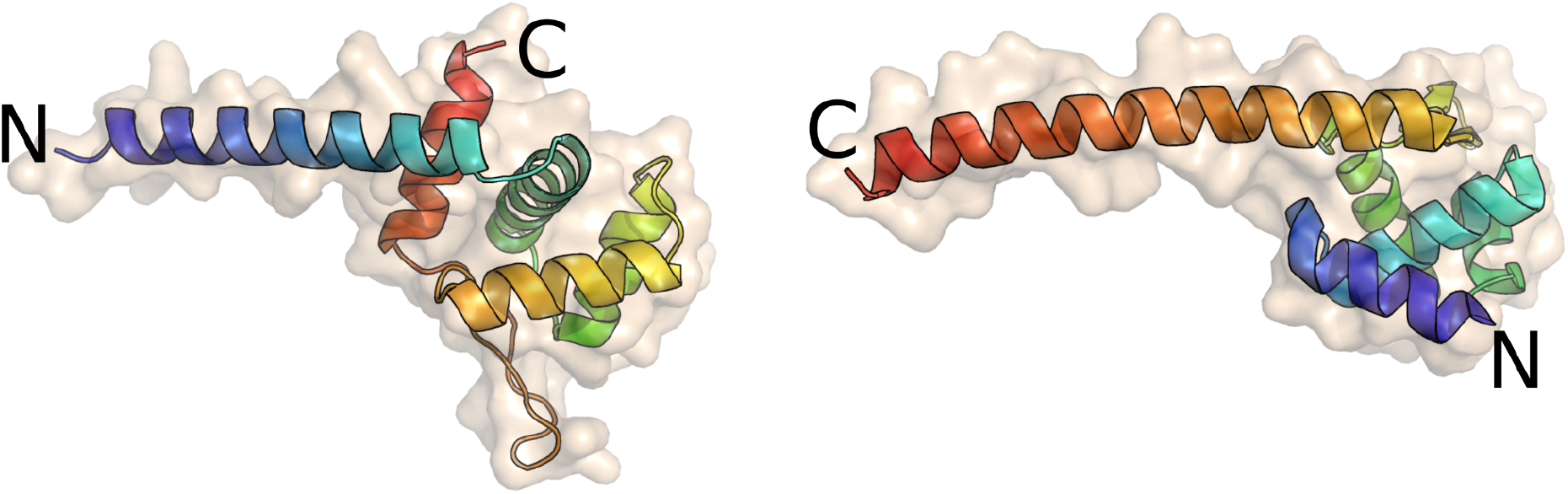
Comparison between an archaeal HTH domain (5box chain A:1-111) and an unannotated Asgard sequence. The image displays a crystal structure of an archaeal HTH DNA-binding protein (left) and the AlphaFold structure of an unannotated Asgard protein (OLS30097, right). Xenusia predicts the latter with high confidence to be a DNA-binding protein. The color gradient represents the amino acid positions in the sequence, transitioning from blue at the beginning (N-terminal) to red at the end (C-terminal). Although the two proteins share a remarkably similar structure, their orientation is reversed, indicating a potential result of convergent evolution. HTH, helix-turn-helix.

### EXPERIMENTAL CHARACTERIZATION OF ASGARD DNA-BINDING PROTEINS

As mentioned earlier, working with Asgard proteins is often challenging due to weak homology links with proteins of known functions. Consequently, transferring annotations can be risky. To address this, we chose three Asgard proteins (OLS23561, OLS17986, and OLS31011) with weak homology connections to known DNA proteins. These proteins are confidently predicted by Xenusia to be DNA-binding proteins. We then conducted experimental investigations on their capability of interacting with DNA. Supplementary Table 2 provides details about these selected proteins.

The three proteins were produced in *E. coli* and purified via His-tag affinity chromatography. The capability of these proteins to bind DNA was evaluated with gel electrophoresis followed by autoradiograph, shown in Figure 3. For each protein, we tested 3 different ^32^P-labeled dsDNA probes, named P1, P2 and P3 (see Supplementary table 3). These probes consist of the promoter regions of the genes encoding the putative DNA-binding proteins and the promoter regions of genes in the proximity of the target gene (see method section DNA-binding assay). We made this choice because in archaea, transcription factors frequently influence genes that are proximate in the sequence space. This assay should provide a first estimation of the binding specificity, as it reveals the migration speed of the protein-dsDNA mixture with respect to the free dsDNA probe. Upon binding, the protein-dsDNA complex will be heavier and therefore migrate slower. Figure 3 clearly shows that OLS17986 binds to all three probes, both in its monomeric form as in an aggregated form. OLS31011 forms only a monomeric complex with its P1 probe. The concentration of OLS23561 appeared to be too high as only aggregated probe-protein complexes could be observed. Therefore, a crescent protein concentration of OLS23561 was assessed to test for monomeric interaction to the probes, as was observed for probes 1 and 2 (see Supplementary Figure 2).

**Figure 3:**
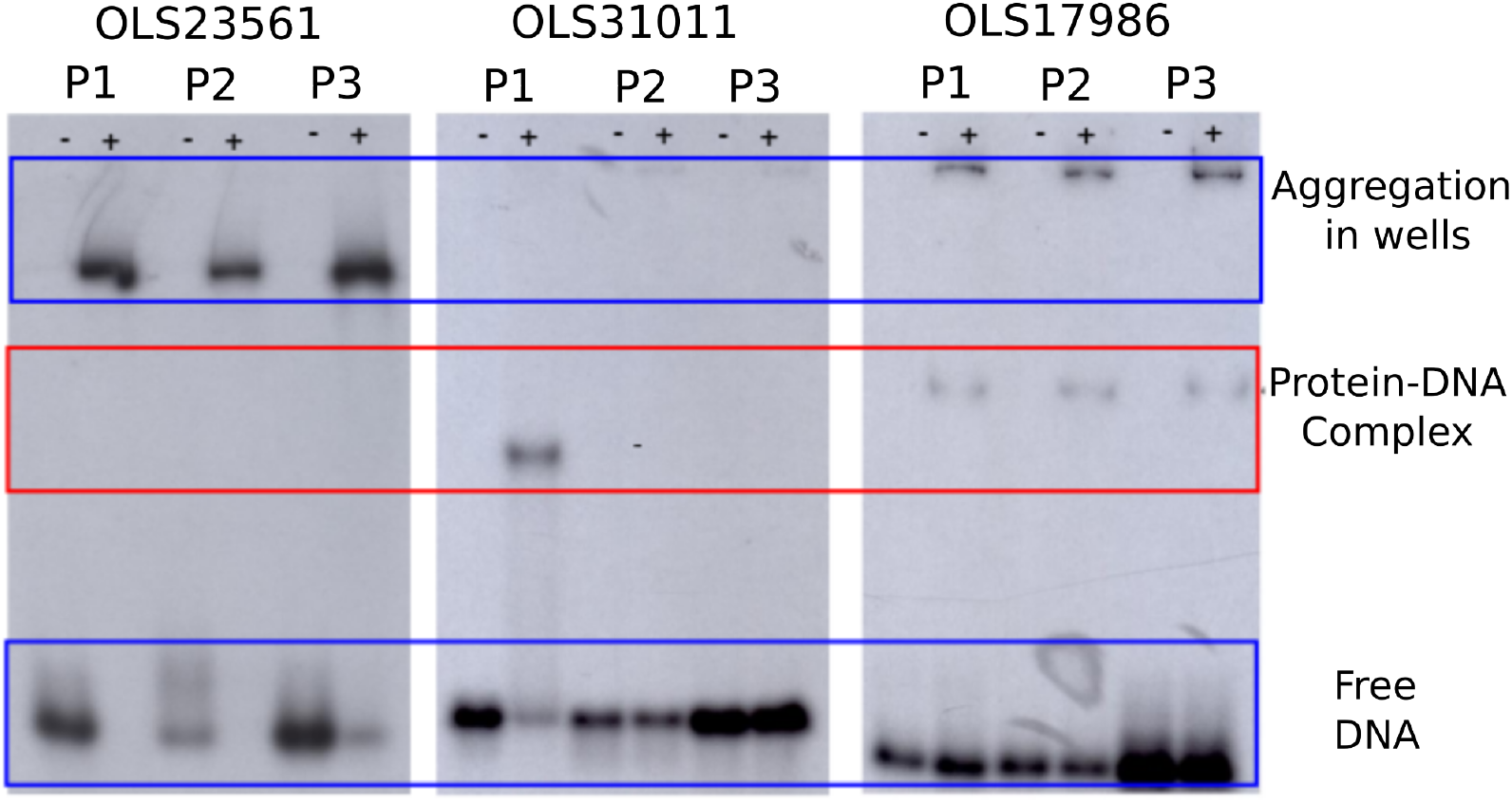
Experimental characterization of three Asgard DNA-binding proteins. The interaction of the three Asgard proteins (OLS23561, OLS31011 and OLS17986) with each three labeled dsDNA probes (P1, P2, and P3) was assessed via gel electrophoresis followed by autoradiography. Wells indicated with a “+” contain protein and DNA, while those indicated with a “-” are the negative control and only contain the DNA probe. OLS23561 does not show monomeric protein-DNA complex formation, only aggregated complexes are detected. OLS31011 only forms a complex with P1, while OLS17986 forms a complex with all three probes. The red box indicates where the band of the monomeric protein-DNA complex should appear.

## METHODS

### DATASETS

For this study, we constructed three distinct datasets from publicly available online resources. Two of these datasets were used exclusively for training and optimization, while the third dataset was reserved for final validation. The training datasets include: (1) the contact dataset, comprising protein sequences derived from PDB structures, where each residue is annotated with a binary label indicating whether the corresponding amino acid interacts with DNA; and (2) the domain dataset, which incorporates per-residue annotations from UniProt regarding DNA-binding domains. To reduce the risk of overfitting, both of these datasets contain only bacterial proteins.

The third dataset, designated as the validation dataset, contains solely archaeal proteins. This dataset was not used for any optimization or hyperparameter tuning to further mitigate overfitting risks. Additionally, all datasets were curated to ensure that both internal sequence identity and cross-dataset sequence identity are below 20% (obtained using CD-HIT ^20^), ensuring minimal redundancy.

#### CONTACT DATASET

To construct this dataset, we utilized the filtering function of the PDB website (2018 version) and selected proteins based on the following criteria: (1) the proteins must be of bacterial origin, (2) the structure must have been resolved using X-ray diffraction, (3) the structure resolution must be < 2 Å, and (4) the structure must contain both protein and DNA co-crystallized.

The selected proteins were then annotated using a PyMOL script, which labeled each residue as interacting (1) if it has at least one atom within 1.5 Å of a DNA atom, or non-interacting (0) otherwise. This process resulted in a dataset of sequences where each residue is annotated as either interacting or non-interacting with DNA. The final dataset comprises 78 annotated sequences. All datasets used in this study are available in the associated Git repository.

#### DOMAIN DATASET

This dataset was constructed using the UniProt filtering system by selecting bacterial proteins that have been manually annotated as DNA-binding proteins. To achieve this, we employed the keyword “DNA-binding” (KW-0238) within the UniProt search engine. UniProt also provides annotations for the regions corresponding to DNA-binding domains, enabling us to categorize the residues of each protein as either part of a DNA-binding domain (1) or not (0). The database consists of 162 DNA-binding proteins.

#### VALIDATION DATASET

This dataset is composed exclusively of archaeal proteins and was manually constructed and curated using data from the literature. Proteins labeled as positive were identified in the literature as having DNA-binding activity. In contrast, proteins labeled as negative were selected based on their extensive annotations in UniProt, with no reported DNA-binding activity or any other function involving interactions with nucleic acids. The final dataset comprises 211 proteins, of which 174 are negative hits and 37 are positive hits.

### PREDICTION OF DNA-INTERACTING RESIDUES (NN1)

Every amino acid in the protein sequence is encoded with 4 features: 1) predicted backbone dynamics and secondary structure propensities for 2) sheet, 3) helix and 4) coil (using DynaMine ^21^ similarly as described in ^22^). The final input is defined as a window of 11 residues with the residue to be predicted as the central one (residue no.6), resulting in 44 features (as we did in other tools ^23^ ^24^). The feature vectors are then used to feed a 2 layers gated recurrent units (GRU) neural network with a hidden size of 7, followed by a max-pooling layer and sigmoid activation.

The neural network was trained on the contact dataset for 300 epochs with learning rate of 0.001, batch of 5 and weight decay of 0.0001, using the ADAM optimizer. The network’s training task was to predict which residues are in contact with DNA.

### PREDICTION OF DNA-BINDING DOMAINS (NN2)

As for the development of NN1, every amino acid in the protein sequence is encoded with 4 features: 1) predicted early folding predictions ^22^, 2) backbone dynamics predictions ^21^, and 3) a one-hot encoding vector defining the amino acid. The encoding scheme is then enriched with the addition of 4) the predicted DNA-interacting residues from NN1, which is concatenated to obtain a feature vector describing every residue. The final input is defined by taking a sliding window of 11 contiguous residues and concatenating their feature vectors, as done in previous works ^21^.

The final inputs are used to feed a feed-forward neural network consisting of 3 layers, with each 30 neurons and rectified linear unit (ReLU) activation. The last layer ends in a single neuron with sigmoid activation, providing the final per-residue prediction.

The neural network was trained on the domain dataset for 300 epochs with learning rate of 0.001, batch of 5 and weight decay of 0.0001, using the ADAM optimizer. The network’s training task was to identify DNA-binding domains, taking the output of NN1 as input.

In order to obtain a single prediction per protein, we performed a sliding window averaging using a window size of 21 residues, and took the maximum average window value of the protein.

### EXPERIMENTAL CHARACTERIZATION OF ASGARD DNA-BINDING PROTEINS

#### PLASMID CONSTRUCTION

The pET24a(+) expression vector was used to produce three transcriptional regulator proteins (OLS23561, OLS17986, and OLS31011). This vector allows IPTG-induced T7 polymerase overexpression through its T7 promoter. It also introduces a C-terminal His-tag, enabling protein purification by affinity chromatography. The synthetic DNA fragments of the chosen Asgard genes were inserted into the vector via a unidirectional restriction-ligation reaction, using the FastDigest restriction enzymes NdeI and XhoI (Thermo Fisher Scientific). The vectors were then transformed into competent DH5α *E. coli* cells and plated. After overnight growth at 37 °C, multiple colonies were pooled to perform a colony PCR reaction for the identification of functional expression vectors. Sequencing of all positive candidates validated the sequences for each construct.

#### PROTEIN EXPRESSION AND PURIFICATION

The validated constructs were then transformed into competent Rosetta(DE3) *E*.*coli* cells. These cells, being derivatives of BL21 E. coli cells that contain additional genes for tRNAs recognizing rare E. coli codons, were explored for potentially accelerated translation of heterologous genes. A single colony preculture was grown overnight at 37 °C, diluted 1:60 and allowed to grow until the OD600 reached 0.6. Protein expression was initiated through IPTG addition and incubated for 3 h at 37 °C. Cells were harvested by centrifugation (10 min at 5,000 rpm), washed in 0.9% NaCl and centrifuged again (10 min at 7,000 rpm). The pellet was then lysed chemically and physically, using lysis buffer and sonication respectively. The lysate was centrifuged (10 min at 9,500 rpm) to obtain the soluble and insoluble protein factions. The presence of the protein of interest in these fractions was assessed using SDS-PAGE. The proteins were purified from the soluble fraction through His-tag affinity chromatography, using an ÄKTA-fast protein liquid chromatography system with a HisTrap column (Thermo Fisher Scientific). The protein was eluted from the column by gradually increasing (40 mM to 500 mM) the imidazole concentration. The identity of the eluted protein was confirmed through SDS-PAGE analysis of selected peak fractions.

#### DNA-BINDING ASSAY

To assess the *in vitro* DNA-binding activity of the purified proteins, a DNA-binding analysis was conducted using three ^32^P-labeled dsDNA probes per protein, each approximately 100 bp in length. These probes were chosen to represent the promoter regions proximal to the regulator gene, as many prokaryotic transcription factors exhibit autoregulation, and their target genes are frequently situated in their immediate genomic vicinity.

For OLS17986, probe 1 corresponds to the promoter region of a gene annotated as a putative citramalate synthase (cimA-1), an enzyme involved in the L-isoleucine biosynthesis pathway. Probe 2 corresponds to the promoter region of OLS17986 itself, allowing the exploration of potential autoregulation. Probe 3 contains the promoter region of an open reading frame housing a molybdenum cofactor guanylyltransferase (mobA), a coenzyme F420:L-glutamate ligase, and a phosphoglycerate mutase.

For OLS31011, probe 1 includes the promoter region of OLS31011 and of a rhaT/eamA-type transporter. The other two probes contain promoter regions of hypothetical genes of unknown function.

For OLS23561, probe 1 includes the promoter of a putative alcohol dehydrogenase gene. Probe 2 contains the promoters of OLS23561 itself and a putative stress response protein. Probe 3 includes the promoter of a putative methyltransferase gene.

To perform the assay, reaction mixtures containing ^32^P-labeled dsDNA probe:protein combinations (20,000 cps:(11.24μM, 5.20μM and 64.22μM for OLS17986, OLS31011 and OLS23561 respectively) and salmon sperm competitor DNA (10 mg/ml) were incubated for 25 min at 37 °C. The mixtures were then separated onto an 8% acrylamide gel (180 V for 10 min followed by 130 V for 2h). After the run, the gel was exposed overnight to an autoradiography gel and developed the next day.

