## supplementary for "In-silico identification of archaeal DNA-binding proteins"

### SUPPLEMENTARY TABLES

***Supplementary Table 1: Predicted Asgard proteins with high confidence as DNA-binding proteins.*** *The table lists the top 10 proteins from the Asgard group, ranked by Xenusia's highest confidence scores for their predicted DNA interactions. ABC, ATP-binding cassette; HTH, helix-turn-helix.*

| # | Xenusia Score | Uniparc ID | Annotation |
|---|---|---|---|
| 1 | 0.957 | OLS30781 | Metalloprotease (homology) |
| 2 | 0.954 | OLS30429 | Characterized HTH |
| 3 | 0.950 | OLS21705 | Characterized HTH |
| 4 | 0.948 | OLS27061 | ABC transporter (homology) |
| 5 | 0.947 | OLS17998 | HTH (homology) |
| 6 | 0.947 | OLS16705 | HTH (homology) |
| 7 | 0.945 | OLS19303 | Response regulator SaeR |
| 8 | 0.945 | OLS28978 | Aminopeptidase BapA (homology) |
| 9 | 0.943 | OLS30097 | - |
| 10 | 0.941 | OLS18037 | S60 ribosomal protein (homology) |

***Supplementary Table 2: Three experimentally investigated Asgard proteins.*** *The table presents the three Asgard proteins that underwent experimental examination, along with their distant homologous transcription factors. The homologous protein and the sequence identity has been identified using the BLAST web server [(Johnson et al. 2008)](#).*

| Xenusia Score | Uniparc ID | Homologous Organism | Sequence Identity |
|---|---|---|---|
| 0.904 | OLS17986 | Sulfurisphaera tokodaii (archaea) | 41% |
| 0.926 | OLS23561 | Bacillus subtilis (bacteria) | 43% |
| 0.895 | OLS31011 | Haloferax volcanii (archaea) | 43% |

***Supplementary Table 3:*** *List of dsDNA probes used in the DNA-binding assay.*

| **Name** | **Sequence (5’ - 3’)** |
|---|---|
| *OLS17986*_Probe1f | tgttattcaaatgtgtgaacctctagtttcgagaactagtattacgtgatgttatgtttacctatttaaactttctgatttggtaaaaaaagttttaaaa |
| *OLS17986*_Probe1r | ttttaaaactttttttaccaaatcagaaagtttaaataggtaaacataacatcacgtaatactagttctcgaaactagaggttcacacatttgaataaca |
| *OLS17986*_Probe2f | attagcatattagctcaccttgaccttgtgagatttatatttttaaatatcactgaataataactaactaatacttaaatttaaaccttccagtattct |
| *OLS17986*_Probe2r | agaatactggaaggtttaaatttaagtattagttagttattattcagtgatatttaaaaatataaatctcacaaggtcaaggtgagctaatatgctaat |
| *OLS17986*_Probe3f | aacataatttttctaacatccatcagggagtccacctagcaaagttaaaatgctagtcgattatactaacttttatttaacattaattagaacgcggtga |
| *OLS17986*_Probe3r | tcaccgcgttctaattaatgttaaataaaagttagtataatcgactagcattttaactttgctaggtggactccctgatggatgttagaaaaattatgtt |
| *OLS31011*_Probe1f | tcaatcaagagattcattctcccaagtaaataggtagattgaagtaggaatagtgttttgccgtcatgcgacgggtgtttataagatgatagaagctaaa |
| *OLS31011*_Probe1r | tttagcttctatcatcttataaacacccgtcgcatgacggcaaaacactattcctacttcaatctacctatttacttgggagaatgaatctcttgattga |
| *OLS31011*_Probe2f | gaatatctctgacaatcatggataacagtgaaatagcattgtgtcatgggtacaatgattacgatggtggattcggagtacttacaggataagaaagttc |
| *OLS31011*_Probe2r | gaactttcttatcctgtaagtactccgaatccaccatcgtaatcattgtacccatgacacaatgctatttcactgttatccatgattgtcagagatattc |
| *OLS31011*_Probe3f | ggttgaggatgaaaagaaaagagagattggacttcgactaacgaattcaatgggaggcttgaggagtgaacgcatatttcacaatagtactacttggatt |
| *OLS31011*_Probe3r | aatccaagtagtactattgtgaaatatgcgttcactcctcaagcctcccattgaattcgttagtcgaagtccaatctctcttttcttttcatcctcaacc |
| *OLS23561*_Probe1f | cacatatatactttttatagtaatattggttacaaatgttttcacctaattaaaaaacaattaataatgatatatttcgaggaatacctatgagagcaat |

| Name | Sequence (5’ - 3’) |
| --- | --- |
| *OLS23561*_Probe1r | attgctctcataggtattcctcgaaatatatcattattaattgttttttaattaggtgaaaacatttgtaaccaatattactataaaaagtatatatgtg |
| *OLS23561*_Probe2f | aatattcatttaaccacatggttgaatataatttagtgtatataaatatttcattcaactaaatggtataatgtttcttatccaatcaaataccaatatc |
| *OLS23561*_Probe2r | gatattggtatttgattggataagaaacattataccatttagttgaatgaaatatttatatacactaaattatattcaaccatgtggttaaatgaatatt |
| *OLS23561*_Probe3f | caaaaaccatatctcaattctcttaaattacttgcattttgcaagagaatcttattctgcatattcttaactaattattaatttggttcttaacttatcg |
| *OLS23561*_Probe3r | cgataagttaagaaccaaattaataattagttaagaatatgcagaataagattctcttgcaaaatgcaagtaatttaagagaattgagatatggtttttg |

### SUPPLEMENTARY FIGURES

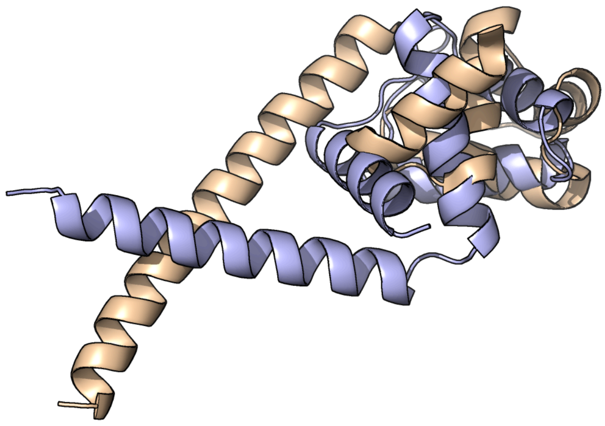

***Supplementary Figure 1: Alignment of the AlphaFold predicted structure of an unannotated Asgard protein and the crystal structure of an archaeal HTH DNA-binding protein.*** *The image shows the structural alignment (performed with PyMOL) of the two protein structures shown in Figure 2. While the structure is similar, their sequence is reversed and the algorithm fails to align them.*

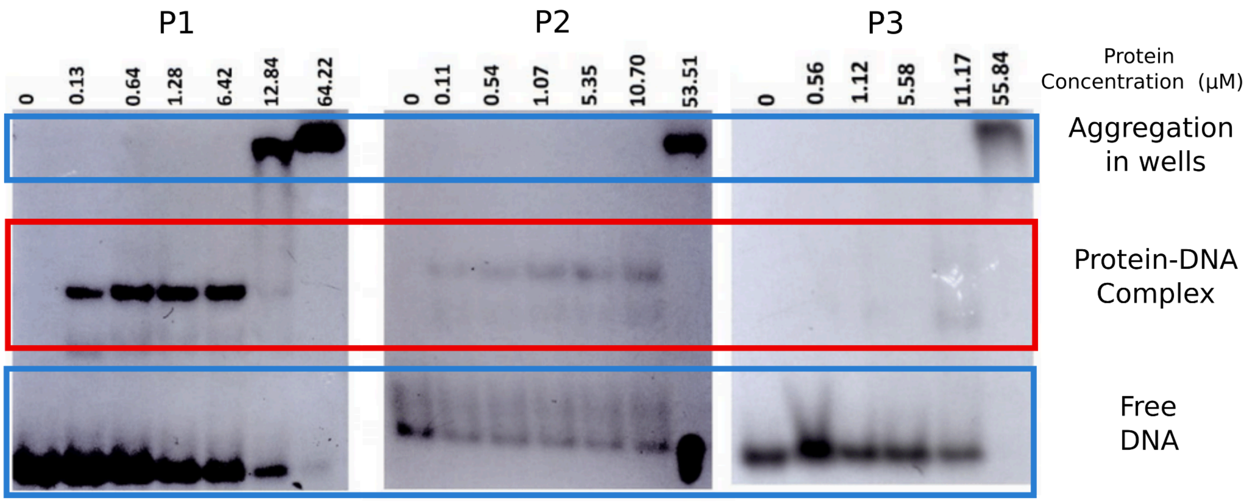


***Supplementary Figure 2: DNA binding assay for OLS23561 using different protein concentrations.*** *The image shows the results of the DNA binding assay for OLS23561 at varying protein concentrations, addressing the aggregation problems observed in Figure 3. At low concentrations (<10 µM), no protein aggregation is observed. The red box indicates where the band of the monomeric protein-DNA complex should appear. The test reveals that the protein binds specifically to probes P1 and P2 but not to P3.*
